# HISAT-genotype: Next Generation Genomic Analysis Platform on a Personal Computer

**DOI:** 10.1101/266197

**Authors:** Daehwan Kim, Joseph Paggi, Steven L. Salzberg

## Abstract

Rapid advances in next-generation sequencing technologies have dramatically changed our ability to perform genome-scale analyses of human genomes. The human reference genome used for most genomic analyses represents only a small number of individuals, limiting its usefulness for genotyping. We designed a novel method, HISAT-genotype, for representing and searching an expanded model of the human reference genome, in which a comprehensive catalogue of known genomic variants and haplotypes is incorporated into the data structure used for searching and alignment. This strategy for representing a population of genomes, along with a very fast and memory-efficient search algorithm, enables more detailed and accurate variant analyses than previous methods. We demonstrate HISAT-genotype’s accuracy for HLA typing, a critical task in human organ transplantation, and for the DNA fingerprinting tests widely used in forensics. In both applications, HISAT-genotype not only improves upon earlier computational methods, but matches or exceeds the accuracy of laboratory-based assays.

**One Sentence Summary:** HISAT-genotype is a software platform that has the ability to genotype all the genes in an individual’s genome within a few hours on a desktop computer.

---

Advancements in sequencing technologies and computational methods have enabled rapid and accurate identification of genetic variants in the human population. The individual genomic data revealed through these advancements along with relevant clinical and environmental information promise to help improve predictions for cancer risk, inform lifestyle choices, generate more accurate clinical diagnoses, reduce adverse drug reactions and other negative side effects of treatments, and improve patient outcomes through better-targeted therapies. Although massive sequencing projects over the past decade such as the 1,000 Genomes Project [1, 2], GTEx [3], GEUVADIS [4, 5], and the Simons Simplex Collection (SSC) [6, 7] have generated trillions of reads that are available from public archives [8], our ability to make use of these enormous data sets is quite limited. One important limitation is that analyses must rely on the alignment of sequencing reads against the human reference genome [9], which does not reflect genetic diversity across individuals and populations. Sequences from other humans, particularly those not included in the samples used for constructing the human reference, may align incorrectly or not at all when they originate from a region that differs from the reference genome. This reliance on a single reference genome can introduce significant biases in downstream analyses, and it can miss important disease-related genetic variants if they occur in regions not present in the reference genome.

A series of large-scale projects in recent years have yielded >110 million SNPs (in dbSNP [10]) and >10 million structural variants (in dbVar [11]). Although these variants represent a valuable resource for genetic analysis, current computational tools do not adequately incorporate them into genetic analysis. For example, >3,000 alleles of the HLA-A gene, which must be matched precisely between donors and recipients of organ and stem cell transplants, have been identified. Representing and searching through the numerous alleles of even one gene has been a challenge requiring a large amount of compute time and memory. Computational methods have thus focused on genotyping one or a few genes because whole-genome genotyping has simply been impractical.

To address these challenges, we have developed a novel indexing scheme that uses a graph-based approach to capture a wide representation of genetic variants with very low memory requirements. We have built a new alignment system, HISAT2 (ccb.jhu.edu/software/hisat2), that enables fast search through the index. HISAT2 is the first and currently the only practical method available for aligning raw sequencing reads to a graph that captures the entire human genome. Our graph-based alignment approach enables much higher alignment sensitivity and accuracy than standard, “linear” reference-based alignment approaches, especially for highly polymorphic genomic regions. Using HISAT2 as a foundation, we developed HISAT-genotype to compute the HLA type and the DNA “fingerprint” of a human using standard whole-genome sequencing data. Because HISAT-genotype works well for multiple highly diverse genes and genomic regions, we expect that it will be straightforward to extend it to many more known variants in human genes. HISAT-genotype is open-source software freely available at http://www.ccb.jhu.edu/software/hisat-genotype.

## Results

To demonstrate the capability of HISAT-genotype, we describe results from two experiments: (1) genotyping the human leukocyte antigen genes (HLA-A, HLA-B, HLA-C, HLA-DQA1, HLA-DQB1, HLA-DRB1), which are among the most diverse human genes; and (2) evaluating DNA fingerprinting loci using 13 markers plus the sex-determining marker gene Amelogenin, which are widely used in criminal forensics to identify individuals.

### HLA typing

The IMGT/HLA Database [12] encompasses >16,000 alleles of the HLA gene family. We built a HISAT2 index of the human genome that incorporates all of these variants, which increased the computational resource requirements only slightly as compared to an index without the variants. For highly polymorphic regions such as those containing the HLA genes, HISAT2 is more sensitive than other short-read aligners; e.g., on one of our data sets, HISAT2 maps 12–100% more reads to the HLA genes than Bowtie2 [13](Table S1).

The HLA allele nomenclature uses a set of four numbers from left to right to designate alleles first classified by (1) allele group according to serological and cellular specificities, then further sub-grouped by (2) protein sequence, and similarly subcategorized according to (3) coding and then (4) noncoding sequences; e.g., HLA-A*01:01:01:01 is a specifier for one allele of the HLA-A gene. HISAT-genotype reports alleles for all four fields, unlike many other programs, which tend to report a subset of the numbers (typically the first two numbers). We conducted computational experiments using Illumina’s Platinum Genomes (PG), which consists of 17 genomes (CEPH pedigree 1463, Supplementary Figure 1) that have been sequenced previously (whole genome sequencing data are available [14], hereafter referred to as PG data). Alleles of HLA-A, HLA-B, and HLA-C for the NA12878, NA12891, and NA12982 genomes have been identified using targeted sequencing [15]. A recent study [16] reported the alleles of all six HLA genes for the 17 genomes by applying several computational methods to the PG data, with the results corresponding to the pedigree. Our experiments show that HISAT-genotype’s results exactly match known alleles and computationally identified alleles of the six genes for the 17 genomes, and that its speed surpasses other currently available methods, primarily due to HISAT-genotype’s alignment engine, HISAT2 (Table S2 and Supplementary File 1).

In addition to identifying alleles for each genome, HISAT-genotype is the first method that can use raw sequence data to assemble and report full-length sequences for both alleles of each of the 6 HLA genes, including exons and introns (Fig 1 and Supplementary Figure 2). The complete sequences of HLA-A reported by HISAT-genotype on the 17 genomes are all in perfect agreement with those previously reported. Its assembled sequences for HLA-B, HLA-C, HLADQA1, and HLA-DQB1 are nearly identical to the previously reported ones. The sequences assembled for HLA-DRB1 are accurate but somewhat fragmented, consisting of a small number of contigs. Greater read lengths should enable HISAT-genotype to produce complete sequences for the HLA-DRB1 gene.

**Fig 1.**
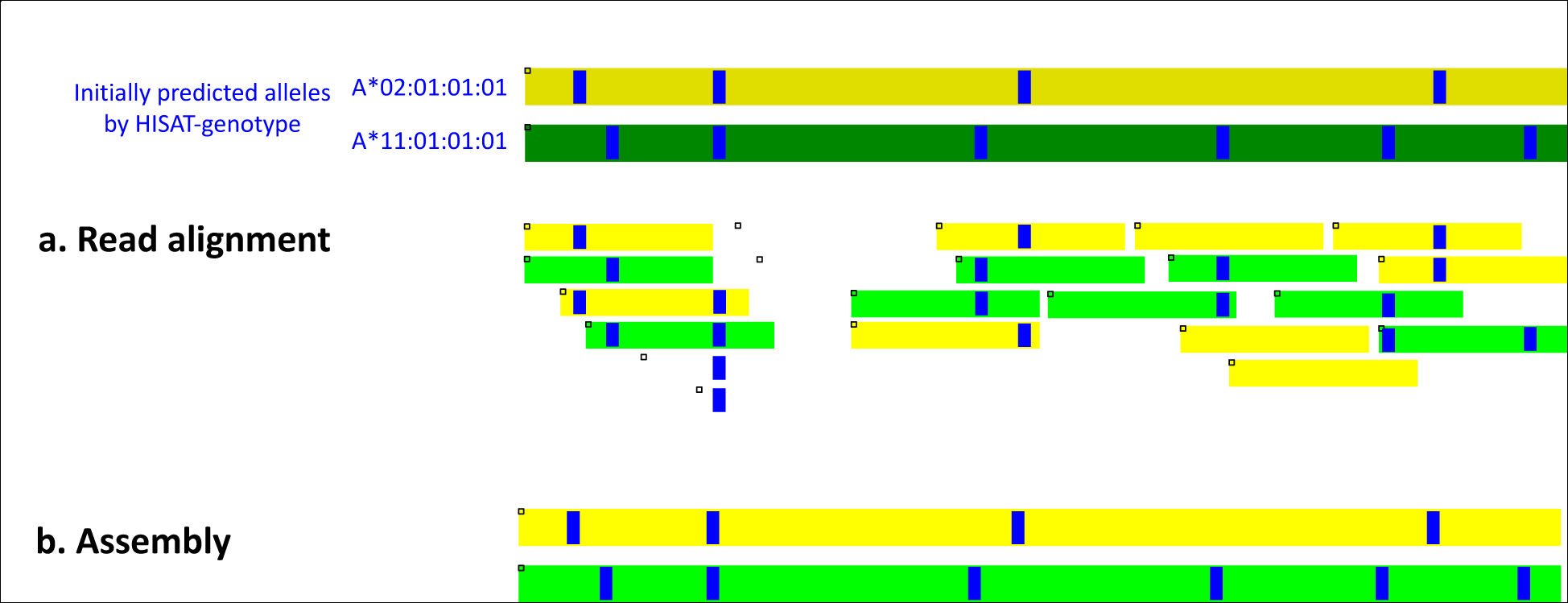
HISAT-genotype’s typing and assembly of HLA genes. The figure shows an abridged example of HISAT-genotype’s assembly output. (Supplementary Figure **2** shows the full assembly output for NA12892, one of the Illumina Platinum genomes.) The two bands shown at the top are the two known alleles predicted by HISAT-genotype, in this case A*02:01:01:01 in dark yellow and A*11:01:01:01 in dark green. Each blue stripe indicates the location of a specific genomic variant with respect to the consensus sequence of the HLA-A gene. *a.* Shorter bands indicating read alignments whose color is determined according to their degree of compatibility with either of the initially predicted alleles. If a read is equally compatible with both alleles, it is shown in white. *b*. Two alleles assembled *de novo* by HISAT-genotype shown in yellow and green, which agree perfectly with the known alleles shown at the top.

In a separate experiment, we compared HISAT-genotype with the Omixon genotyping system [17], an established commercial platform, using whole genome sequencing (WGS) data from the Consortium on Asthma among African-ancestry Populations in the Americas (CAAPA) [18] (Supplementary File 2). Table 1 shows a high concordance rate between the two methods for the allele group and protein sequences (the first two numbers of the HLA classification); more specifically, a concordance of ≥0.97 for genotyping of HLA-A, HLA-B, HLA-C, and HLA-DQA1; 0.91 for HLA-DQB1; and 0.87 for HLA-DRB1. Tests using the CAAPA data also revealed a handful of novel sequences of HLA-A and other HLA genes (Fig 2 and Supplementary Figure 3).

**Fig 2.**
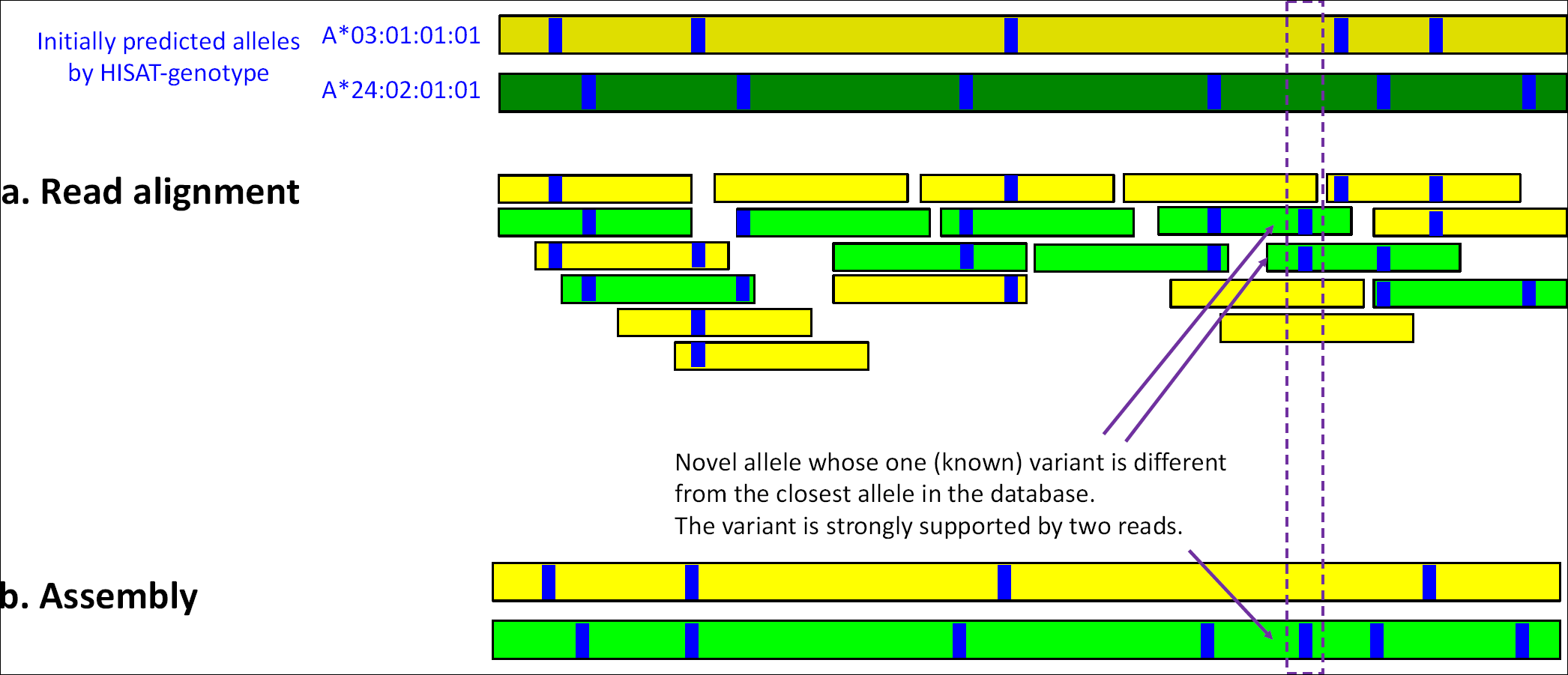
A novel HLA-A allele identified with strong computational evidence. This figure shows an abridged example of HISAT-genotype’s assembly output. At the top are shown the two initially predicted alleles, which are the best matches of the data to previously-known HLA-A alleles. The green assembled allele at the bottom, which was generated *de novo* by HISAT-genotype’s assembler, has one variant different from the predicted allele, A*24:02:01:01. Two reads shown in green support the variant. See Supplementary Figure **3** for more detailed output from a similar case found in LP6005093-DNA_E03 (a CAAPA genome) at the 2,780^th^ base.

**Table 1.** Concordance between HISAT-genotype and Omixon on HLA-typing of 917 genomes from the CAAPA (Consortium on Asthma among African-ancestry Populations in the Americas) collection. Concordance is calculated as the total number of alleles matched between both programs divided by the total number of alleles. For example, for the HLA-A gene, HISAT-genotype and Omixon agree on the allele group (the first number of the HLA type) for both alleles for 913 genomes, agree on one allele for 4 genomes, and agree on no alleles for 0 genomes. Thus, the concordance for HLA-A is 0.998 = (913 × 2 + 4) / (917 × 2). HISAT-genotype reports HLA types with all four fields specified (e.g., A*24:02:01:01), while Omixon reports HLA types with either two numbers (e.g. A*69:01) or three numbers (A*24:02:01); therefore matches were evaluated using only the first two numbers.

|  | First number (e.g., A*01) |  |  |  | First and second numbers (e.g., A*01:01) |  |  |  |
| --- | --- | --- | --- | --- | --- | --- | --- | --- |
|  | Both alleles matched | One allele matched | No allele matched | Concordance | Both alleles matched | One allele matched | No allele matched | Concordance |
| HLA-A | 913 | 4 | 0 | 0.998 | 883 | 33 | 1 | 0.981 |
| HLA-B | 911 | 6 | 0 | 0.997 | 877 | 40 | 0 | 0.978 |
| HLA-C | 915 | 2 | 0 | 0.999 | 880 | 34 | 3 | 0.978 |
| HLA-DQA1 | 884 | 33 | 0 | 0.982 | 868 | 45 | 4 | 0.971 |
| HLA-DQB1 | 917 | 0 | 0 | 1 | 753 | 164 | 0 | 0.911 |
| HLA-DRB1 | 861 | 56 | 0 | 0.97 | 698 | 205 | 14 | 0.873 |

### DNA fingerprinting

DNA fingerprinting analysis has been widely used in criminal investigations and paternity testing since its introduction in the mid-1980’s. It considers a set of 13 highly polymorphic regions that in combination can uniquely identify individuals or their close relatives. The billions of reads in a whole-genome sequencing run include those from the 13 genomic regions used for DNA fingerprinting analysis. In addition to running HISAT-genotype on the WGS data, we performed traditional wet-lab based DNA-fingerprinting using DNA samples of the 17 PG genomes (Epstein-Barr virus transformed B-lymphocytes), which were purchased from the Coriell Institute, and a DNA fingerprinting kit, PowerPlex^®^ Fusion System from Promega.

HISAT-genotype’s initial results for the PG data almost perfectly match our wet-lab results for 11 out of 13 DNA fingerprinting loci on all 17 genomes and correctly determines sex (using the Amelogenin locus) for all 17 genomes (Supplementary File 3 and Supplementary File 4). In order to identify the potential sources of the discrepancies for the two loci, we examined PG’s raw sequencing data and found that the NIST database used by HISAT-genotype (Supplementary File 5) was missing some alleles of the 17 PG genomes (Supplementary File 6). After incorporating the missing alleles, HISAT-genotype’s results perfectly match the wetlab results for all but 8 cases, which are highlighted in yellow in Supplementary File 7.

Assuming there are no germline and somatic mutations in the PG cell lines, an analysis of the 8 disagreements indicates that HISAT-genotype is correct in all 8 cases. For example, on genome NA12886 at locus D5S818, HISAT-genotype reports two alleles 10 and 12, and the wet-lab method reports three alleles 9, 10, and 12. The pedigree information (Supplementary Figure 1) shows that NA12886’s father (NA12877) has two alleles 10 and 11, and the mother (NA12878) has homozygous allele 12, suggests that allele 9 detected by the wet-lab method is likely a false positive. Another example is NA12877’s D3S1358 locus, for which HISAT-genotype gives more specific results that consist of two different alleles 16 and 16’, which are of the same length but are slightly different in their sequences (allele 16: TCAT followed by three repeats of TCTG, then followed by twelve repeats of TCTA; and allele 16’: TCAT followed by two repeats of TCTG, then followed by thirteen repeats of TCTA). Because the two alleles have identical lengths, the wet-lab method cannot distinguish them and reports just one allele.

The current implementation of HISAT-genotype requires exact sequences of alleles, though this requirement can be somewhat relaxed when performing DNA fingerprinting. On the other hand, knowledge of exact sequences allows us to identify alleles more specifically at base-level resolution.

## Algorithmic details

Here we describe the algorithms underlying HISAT2 and HISAT-genotype. HISAT2 implements a novel graph-based data structure along with an alignment algorithm to enable fast and sensitive alignment of sequencing reads to a genome and a large collection of small variants. HISAT-genotype uses HISAT2 as an alignment engine along with additional algorithms to perform HLA-typing and DNA fingerprinting analysis.

## Graph representation of human populations and alignment (HISAT2)

The reference human genome currently used by most researchers was assembled from data representing only a few individuals, with over 70% of the reference genome sequence coming from only one person [9]. By its very design, the reference does not include genomic variants from the human population. Sequence alignment protocols based on this single reference genome are sometimes unable to align reads correctly, especially when the source genome is relatively distant from the reference genome. HISAT2 begins by creating a linear graph of the reference genome, and then adds insertions, deletions, and mutations as alternative paths through the graph. Fig 3 illustrates how variants are incorporated using a very short reference sequence, GAGCTG. In the graph representation, bases are represented as nodes and their relationships are represented as edges. The figure shows three variants: a single nucleotide polymorphism where T replaces A, a deletion of a T, and an insertion of an A. Although the example shows only 1-base polymorphisms, insertions of up to 20 bps and deletions of any length can be incorporated.

**Fig 3.**
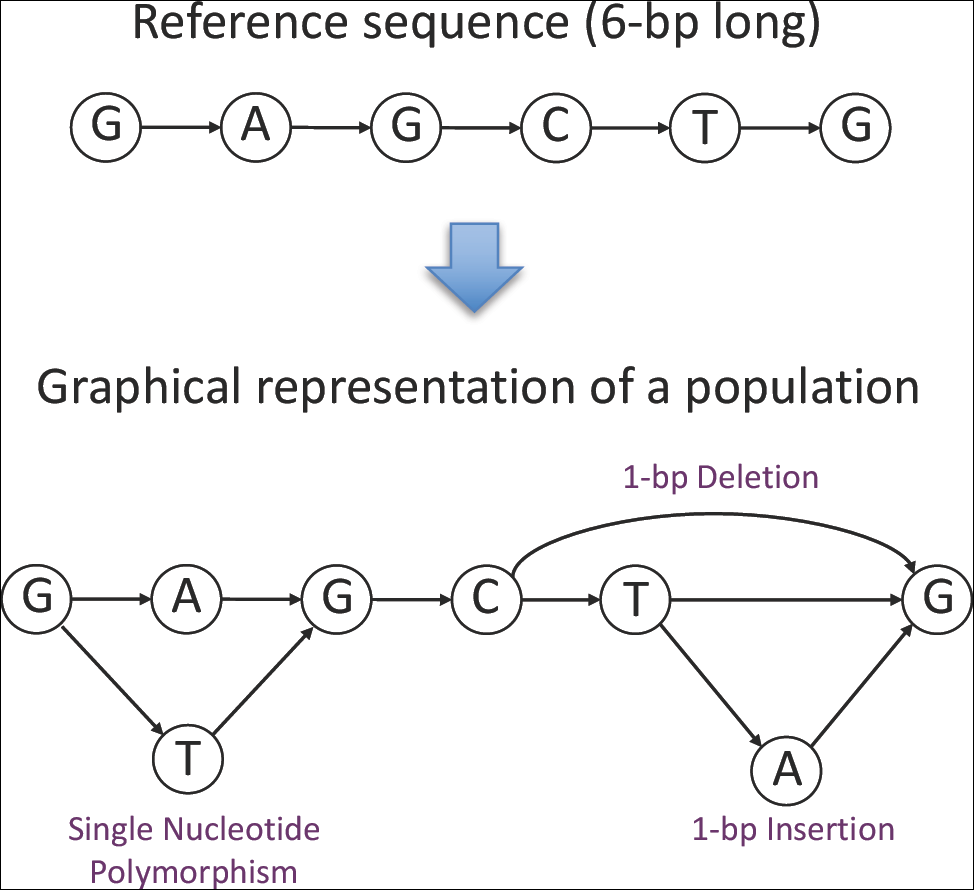
Graph representation of indels and mutation. Starting with a 6-bp reference sequence, GAGCTG (top), the lower graph incorporates three variants: a single nucleotide variant (A/T), a 1-bp deletion (T), and a 1-bp insertion (A).

Any path in the graph defines a string of bases that occur in the reference genome or one of its variants. For example, the path G -> A -> G -> C defines the string GAGC. Strings can be ordered lexicographically; e.g., AGC comes before GTG, which comes before TGZ. A special symbol, Z, is used to indicate the end of the graph and to properly sort strings. To allow fast alignment of queries (reads) to the genome graph, we first convert the graph into a *prefix-sorted graph* using a method developed by Sirén et al [19]. This prefix-sorted graph is more appropriate for search and storage. The prefix-sorted graph is equivalent to the original one in the sense that they define the same set of strings. In a prefix-sorted graph, nodes are sorted such that any strings from a node with a higher lexicographic rank appear before any strings from a node with a lower rank. For example, any string from the node ranked first (node A in Fig 4), such as AGCTGZ, comes before any strings from any other nodes. An equivalent table for this prefix-sorted graph is shown in Fig 5. The table stores two types of information. For outgoing edges, given node rankings 1 to 11, the label of each node is stored according to the number of outgoing edges it has. Here node rankings as also referred to as node IDs. For example, node 1 has one outgoing edge, from A to G, so this node’s label A is stored once, as shown in the first row under “First” of the “Outgoing edge(s)” columns. Node 3 has three outgoing edges, so this node’s label C is stored 3 times. For incoming edges, given the node rankings, the labels of the preceding nodes are stored. For example, node 1 has one incoming edge from the node labeled G, so this G is stored once, in the first row under “Last” of the “Incoming edge(s)” columns. Node 5 has two incoming edges from nodes labeled A and T, so A and T are stored accordingly.

**Fig 4.**
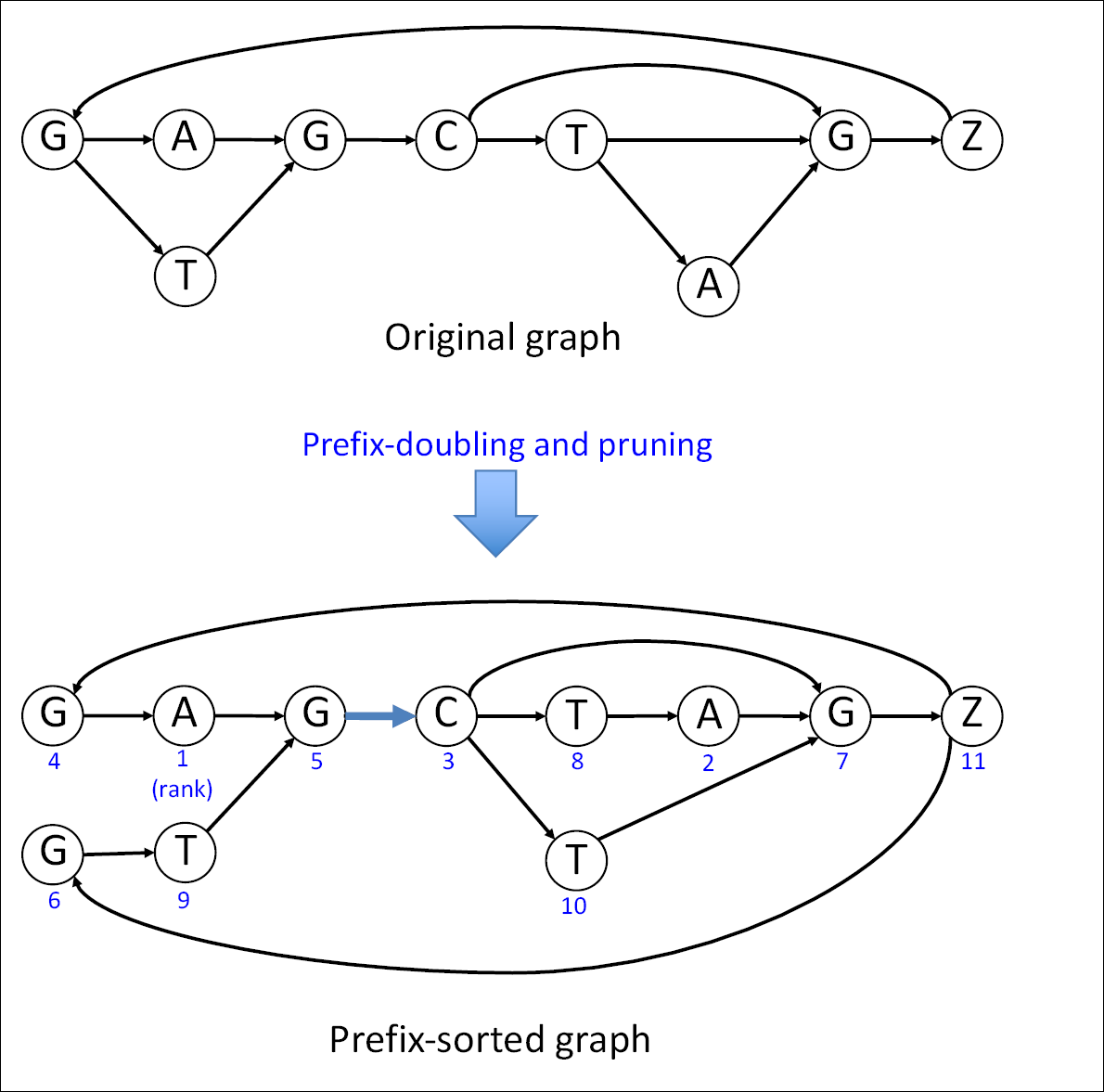
Prefix-sorting the graph. The original graph, with 9 nodes and 12 edges, is augmented so that prefixes can be sorted. The augmented graph has 11 nodes and 14 edges. Each node has a unique numerical node ID shown in blue to indicate its lexicographical order (1 being the first) with respect to the other nodes in the graph. The node labeled with ‘Z’ demarcates the end of the reference sequence.

**Fig 5.**
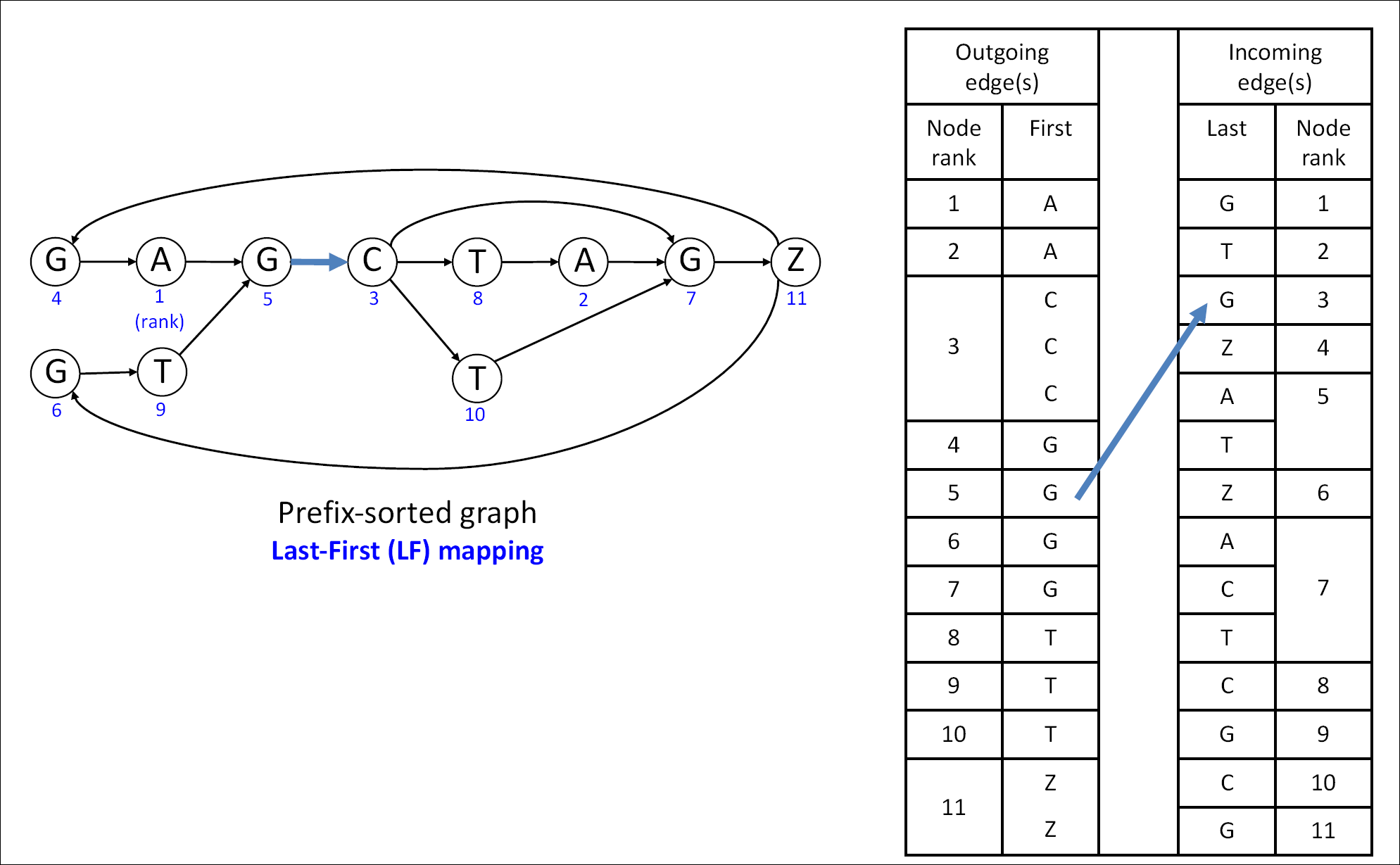
A prefix-sorted graph and its tabular representation. The graph on the left shows 11 nodes and 14 edges. Each node has a unique numerical node ID shown in blue to indicate its lexicographical order (1 being the first) with respect to the other nodes in the graph. The table on the right has two columns under Outgoing edge(s) that show the node IDs and their labels repeated according to the number of their outgoing edges (i.e. node 3, labeled C, is repeated three times with 3 outgoing edges to nodes 7, 8, and 10, respectively). The table has two columns under Incoming edge(s) that show the node IDs and the 14 labels for the preceding nodes (i.e. G is the preceding label for node 1, A and T for node 5). The table is more compact in memory usage than the graph representation.

Although edges are not directly stored using node IDs as depicted in Fig 5, we can implicitly construct the edge information using a very important property of the table representation, called Last-First (LF) mapping. The Last-First mapping property says that the i^th^ occurrence of a certain label in the **last** column corresponds to the i^th^ occurrence of that label in the **first** column. For example, Node 3 in Fig 5 has an incoming edge from the node labeled G. This is the second occurrence of G in the **last** column of the table, which corresponds to node 5 in the **first** column. This indirect representation of edges leads to a substantial reduction in storing the table.

The table representation can be further compacted using the scheme illustrated in Fig 6.

**Fig 6.**
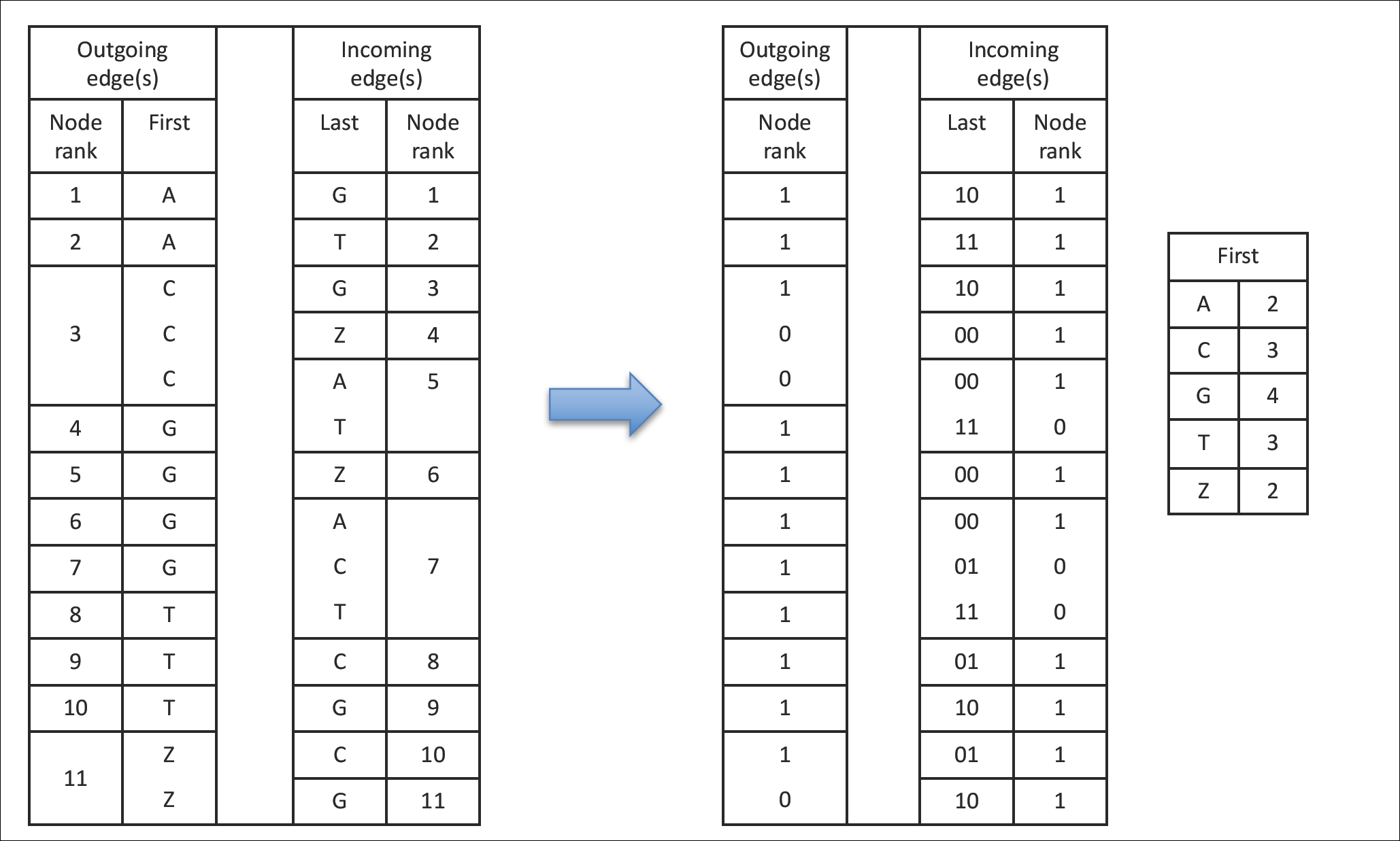
Space efficent representation of the table in Fig 5. In the two ‘Node rank’ columns on the left, since node ranks are given in consecutive and increasing order, one bit (0 or 1) can be used to represent a node rank instead of 4 bytes (any number between 0 and 4,294,967,295) to manage offsets for a human genome. 1 and 0 are used to indicate a new node rank and to indicate an additional outgoing or incoming edge that a node has, respectively. To retrieve a node rank, simply summing up the 1s gives rise to that node’s rank. Since the labels in the ‘First’ column are already sorted, five numbers are enough to represent the column, 2 for As, 3 for Cs, 4 for Gs, 3 for Ts, and 2 for Zs. In the ‘Last’ column, two bits are used to represent each label: 00 for A, 01 for C, 10 for G, and 11 for T. 00 is also used to represent Z. HISAT2 internally resolves whether 00 represents A or Z. The right table is the space efficient representation of the left table after these transformations.

In order to perform the LF mapping, the number of times that a “Last” column label of a given row *r* occurs up to and including *r* needs to be identified, which involves counting occurrences from the top of the table down to row *r*. This counting would be prohibitively time-consuming for the 3-Gb human genome. To accelerate the process, the table is partitioned into small blocks of only hundreds of rows. Additional numbers are stored within each block recording the number of occurrences of a specific base that appear up to that block. We also optimized the local counting process. This overall indexing scheme is called a Graph FM index (GFM) (Fig 7).

**Fig 7.**
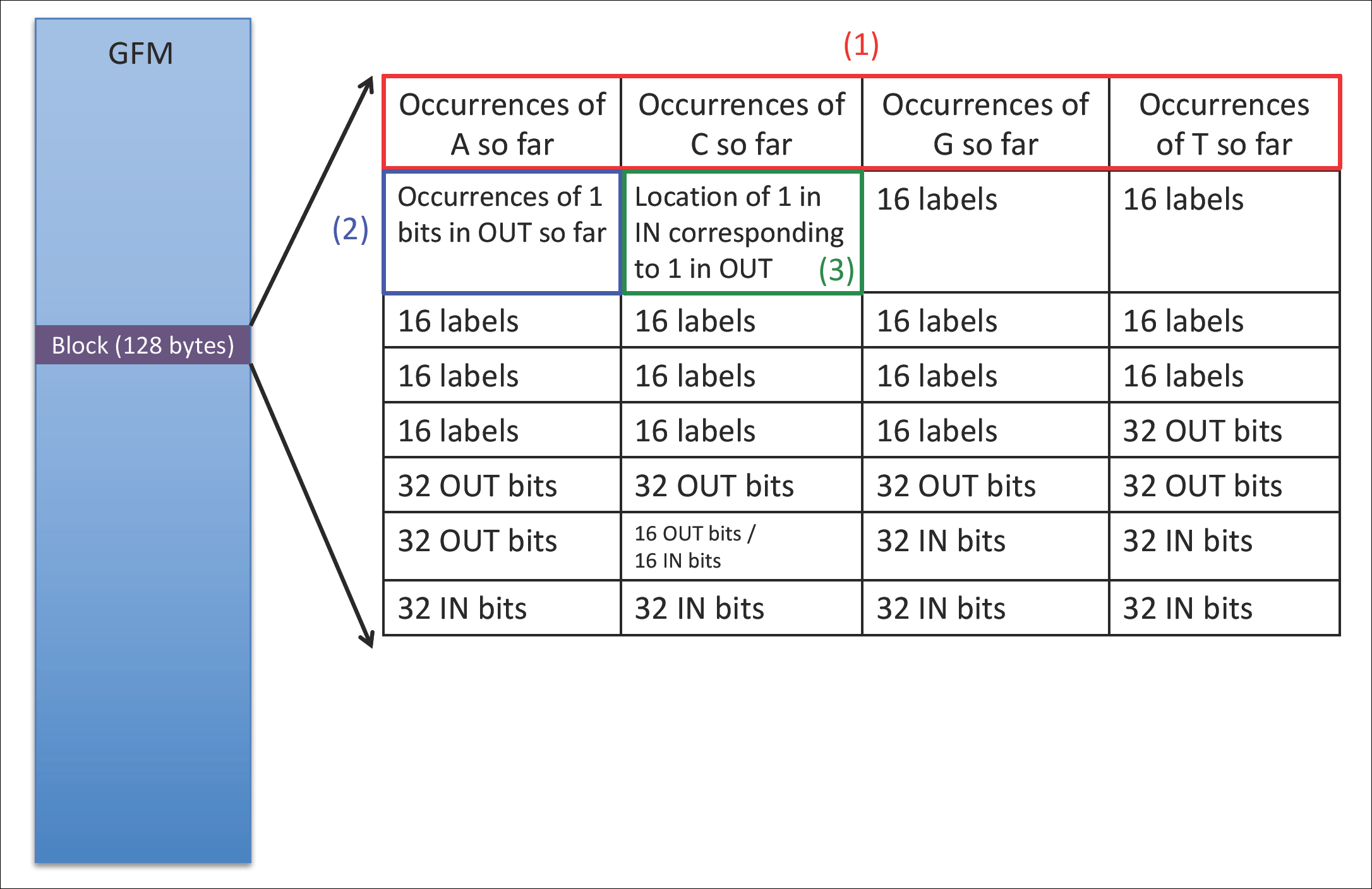
Graph FM index (GFM) The size of each block is 128 bytes consisting of 32 4-byte cells. Each block stores: (1) four 4-byte numbers for the accumulated numbers of occurrences of A, C, G, and T up to that block, (2) one 4-byte number for the accumulated number of 1s up to the block in the Node rank (Outgoing edges) of the right table in Fig 6, (3) one 4-byte number for the row number of the Node rank (incoming edge) corresponding to the accumulated number of 1s indicated in (2), (4) 208 labels (or nucleotides) corresponding to the Last column of the right table in Fig 6, and (5) 208 ‘OUT’ bits and 208 ‘IN’ bits corresponding to the Node rank columns of the right table in Fig 6.

Fig 8 illustrates how a query that contains a known one-base insertion is aligned to the genome using a GFM.

**Fig 8.**
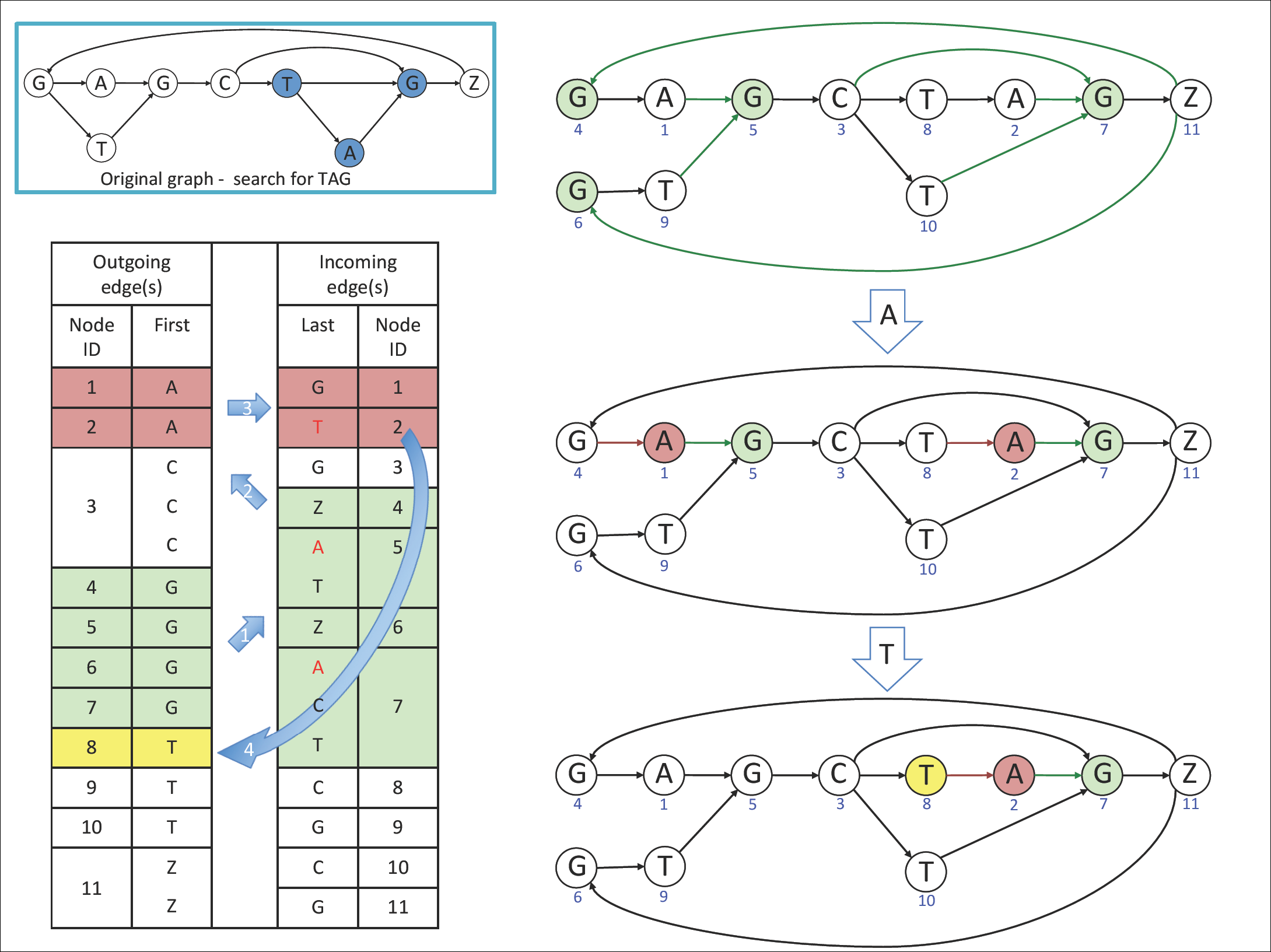
Alignment of a small 3-bp query using a graph FM index. The figure illustrates how to align a 3-bp query, TAG, whose TG corresponds to the last two nucleotides of the original reference sequence, GAGCTG, and A is a 1-bp insertion as shown in Fig 3. Searching from the right end of the query to the left, the nodes labeled ‘G’ are first selected (node IDs ‘4’, ‘5’, ‘6’, and ‘7’). Then the incoming edges of those nodes are examined to identify which has a preceding base ‘A’. Nodes ‘5’ and ‘7’ qualify, with preceding nodes ‘1’ and ‘2’. These in turn are examined to determine which of these nodes is preceded by a base ‘T’. Only one of the two nodes, node ‘2’, has a preceding node, ‘8’, whose label corresponds to ‘T’. Node ‘8’ is chosen as a mapped location for the query. This is the final alignment of the query shown in the prefix-sorted graph, and additional algorithms convert it to the corresponding alignment in the original graph.

To further improve both speed and accuracy, we modified the hierarchical indexing scheme from HISAT [20] to create a Hierarchical Graph FM index (HGFM). In addition to the global index for representing the human genome plus a large collection of variants, we built thousands of small indexes, each spanning ∼57 Kb, which collectively cover the reference genome and its variants. This approach provides two main advantages: (1) it allows search on a local genomic region (57,344 bps), which is particularly useful for aligning RNA-seq reads spanning multiple exons, and (2) it provides a much faster lookup compared to a search using the much larger global index, due to the local index’s small size. In particular, these local indexes are so small that they can fit within a CPU’s cache memory, which is significantly faster than standard RAM.

Our implementation of this new scheme uses just 6.2 GB for and index that represents the entire human genome plus ∼14.5 million common small variants, which include ∼1.5 million insertions and deletions available from dbSNP. The incorporation of these variants requires only 60∼80% additional CPU time compared to HISAT2 (among the fastest alignment programs) searching the human genome without variants, and it obtains greater alignment accuracy for reads containing SNPs (Tables S3 and S4).

## Identification of sequences of genes and genomic regions (HISAT-genotype)

Building off of HISAT2, the next step is to design a graph representation by incorporating known genomic variations and to perform genotyping on a sequencing data set. There is currently no centralized database for the many known genomic variants in human populations. Instead, each database has its own data format and naming conventions. To address this challenge, we parsed exterior databases for human genes or genomic regions and converted them into an intermediate format upon which several HISAT-genotype algorithms are conveniently built. We created a graph genome, called a *Genotype* genome, which is specifically designed to aid in carrying out genotyping as illustrated in Fig 10. In addition to variants and haplotypes, the genotype genome includes some additional sequences inside the consensus sequence shown in yellow, resulting in substantial differences in coordinates with respect to the human reference genome. Thus, it is important to note that a *Genotype* genome should not be used for purposes other than genotyping analysis.

**Fig 9.**
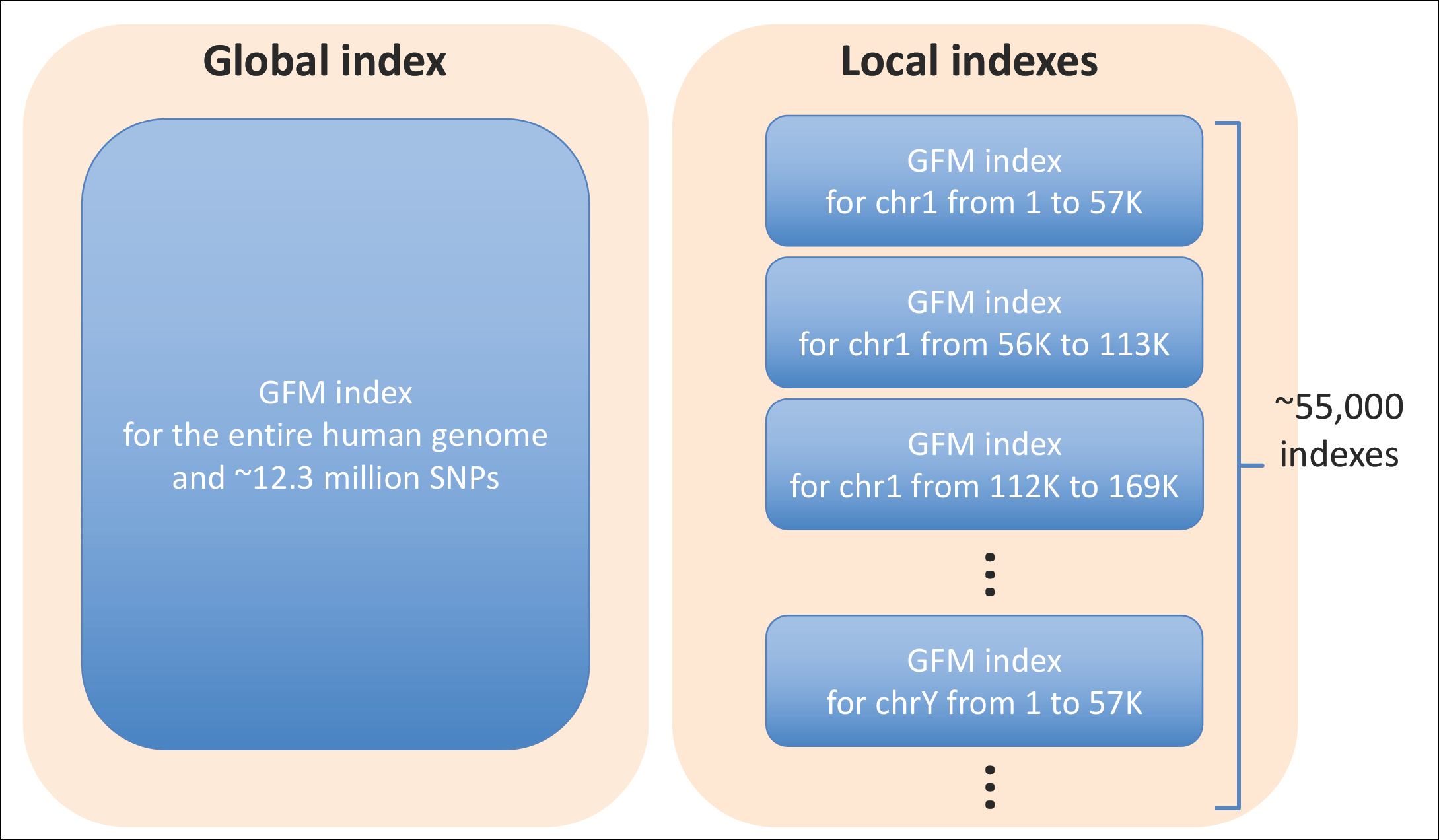
Hierarchical Indexing, i.e., Hierarchical Graph FM index (HGFM) Hierarchical indexing consists of two types of indexes: (1) a global index that represents the entire human genome and (2) 55,172 overlapping local indexes that collectively cover the population. When both are Graph FM-indexes, a genome plus a large collection of variants can be searched simultaneously.

**Fig 10.**
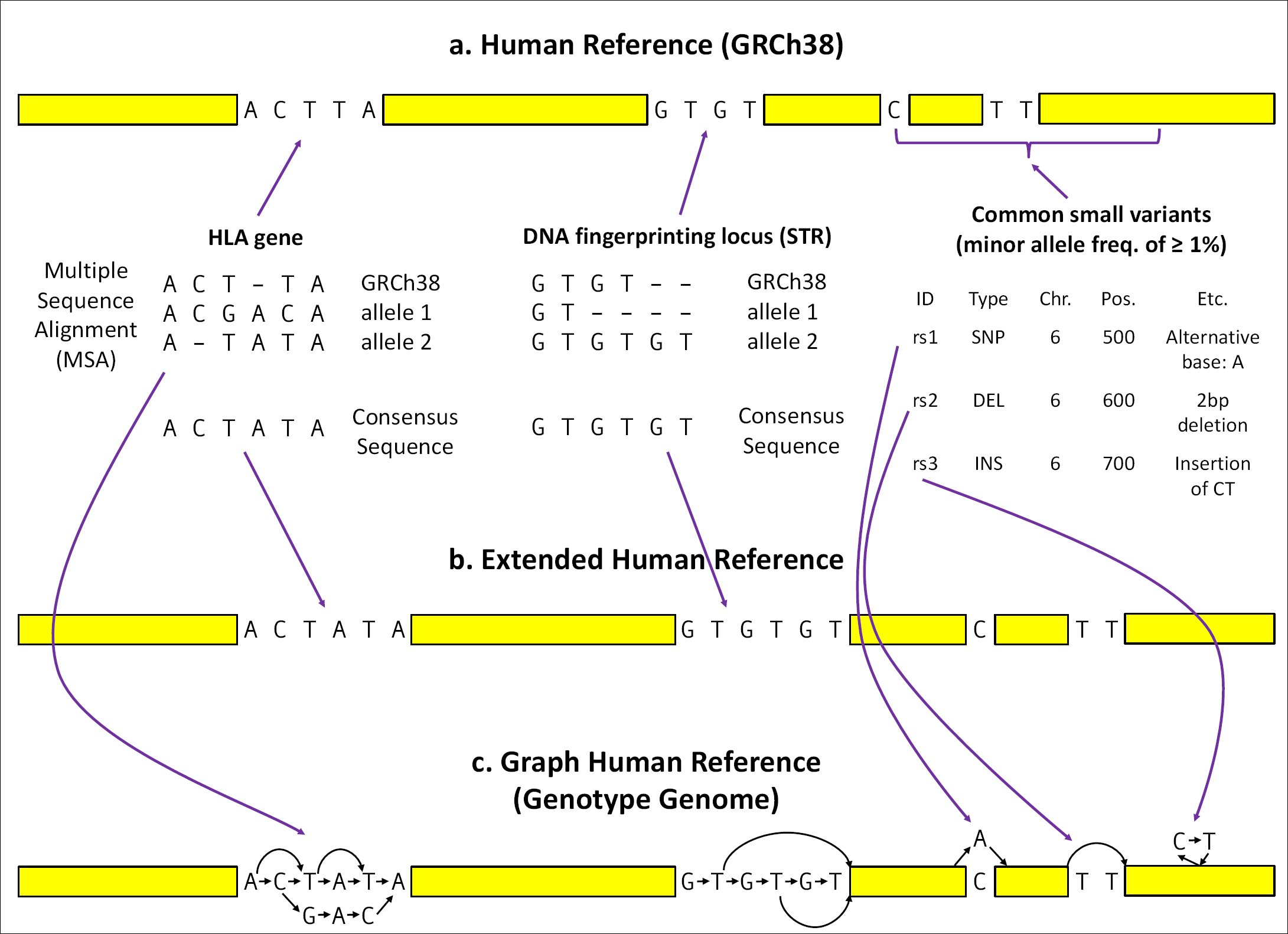
Construction of the Graph Human Reference, i.e. a *Genotype* Genome. The figure illustrates how HISAT-genotype extends the human reference genome (GRCh38) by incorporating known genomic variants from several well-studied genes, DNA fingerprinting loci, and common small variants (i.e. variants with minor allele frequencies of ≥1%) from the dbSNP database. In *a*, the process begins with analyzing information found in the selected databases to construct consensus sequences. The IMGT/HLA database includes over 15,500 allele sequences for 26 HLA genes. A consensus sequence for each HLA gene is constructed based on the most frequent bases that occur in each position of the multiple sequence alignments. The NIST STRBase database contains allele sequences for 13 DNA fingerprinting loci. Because the sequences of the 13 loci are short tandem repeats, HISAT-genotype chooses the longest allele for each locus as a consensus sequence. In *b*, the human reference is extended by replacing the HLA genes and 13 DNA fingerprinting loci with their consensus sequences. In *c*, the known genomic variants are then incorporated into the extended references using nodes and edges. Common small variants from dbSNP such as single nucleotide polymorphisms, deletions, and insertions, are also incorporated into the extended reference. In HISAT-genotype this graph reference is called a *Genotype* genome, upon which a HISAT2 index is built.

In contrast to linear-based representations of the human reference augmented by sequences representing gene alleles, graph representations are much more efficient in terms of memory usage and/or alignment speed, as illustrated in Fig 11. When working with whole-genome sequencing data, using the right reference/index is crucial. Much greater alignment accuracy can be achieved by using a reference that most closely matches the genome where reads were likely to originate. Using the wrong reference (e.g. just a few genes instead of the whole genome) can lead to reads being incorrectly aligned, as depicted in Fig 12.

**Fig 11.**
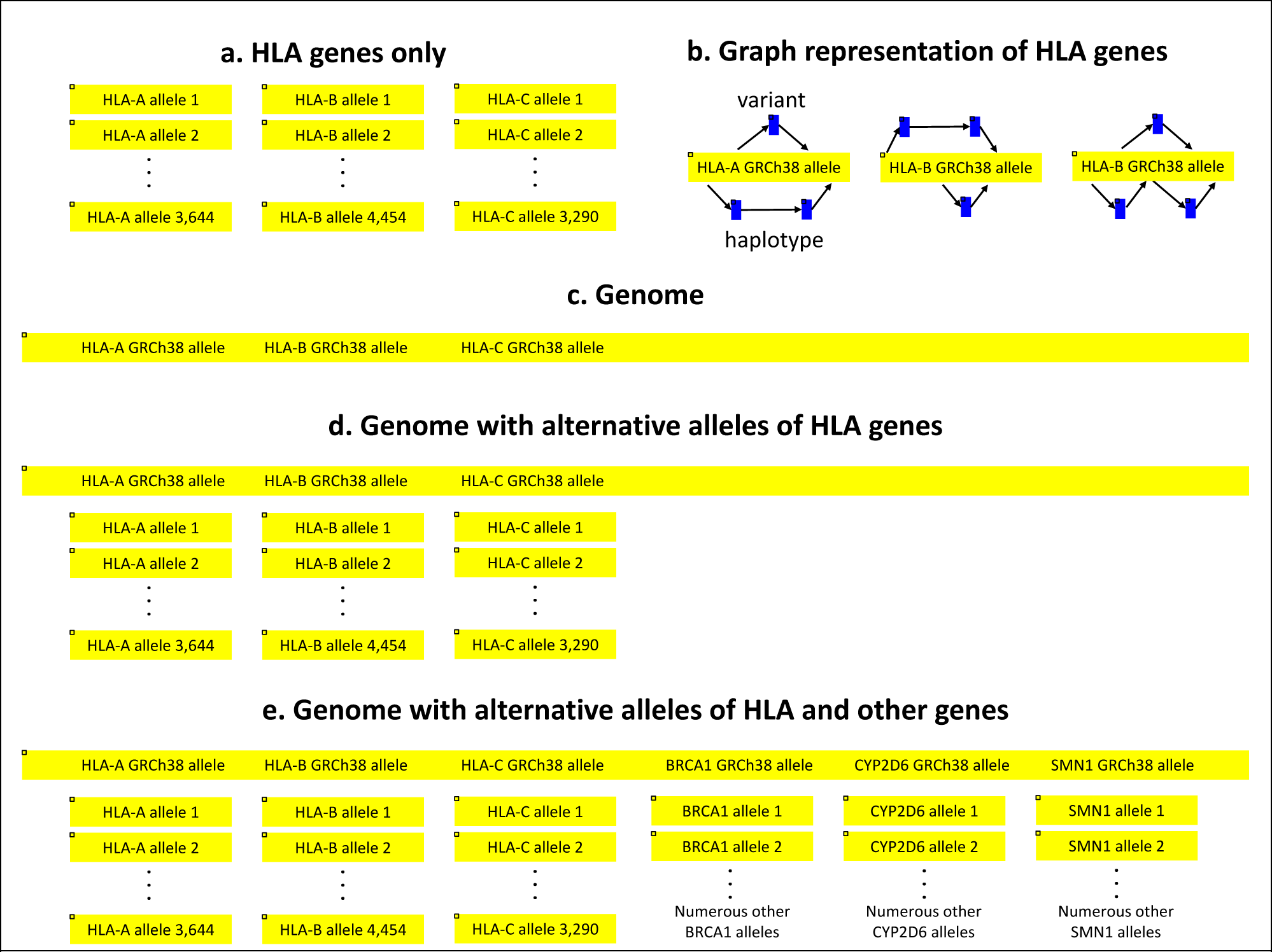
Alternative yet limited approaches to graph representation. Linear reference based representations of many alleles of genes are quite limited in terms of memory usage and/or alignment speed, and have the issue of increased mapping ambiguity. In *a* and *b*, a reference consisting of only alleles of genes of interest can introduce significant mapping bias by mapping reads from regions not included in the restricted reference, as illustrated in more detail in Fig 12. In *c*, the current human reference may not be able to map many reads if they originated from alleles that are substantially different from the human reference allele. In *d*, a reference consisting of the human reference and numerous alleles of HLA genes enables mapping of reads from even substantially different alleles. Most of the HLA-typing methods if not all, such as HLA-VBSeq [21], HLA*PRG [22], Kourami [16], and Graphtyper [23], are based on *c*, *d*, or a combination thereof in order to identify HLA gene reads, after which HLA-VBSeq uses an *a* approach, and HLA*PRG, Kourami, and Graphtyper use a small-scale graph representation as descibed in *b* to perform typing. Kourami assembles only exons of HLA genes, while HISAT-genotype is able to assemble full-length sequences of HLA genes including exons and introns. However, including this many additional alleles introduces considerable mapping ambiguity as most of the alleles are nearly identical to one another, and as a result reads are mapped to many alleles. This approach tends to work on a small number of genes or genomic regions, otherwise, as illustrated in *e*, including alleles of all known genes and genomic regions would require a tremendous amount of memory and greatly increase mapping ambiguity.

**Fig 12.**
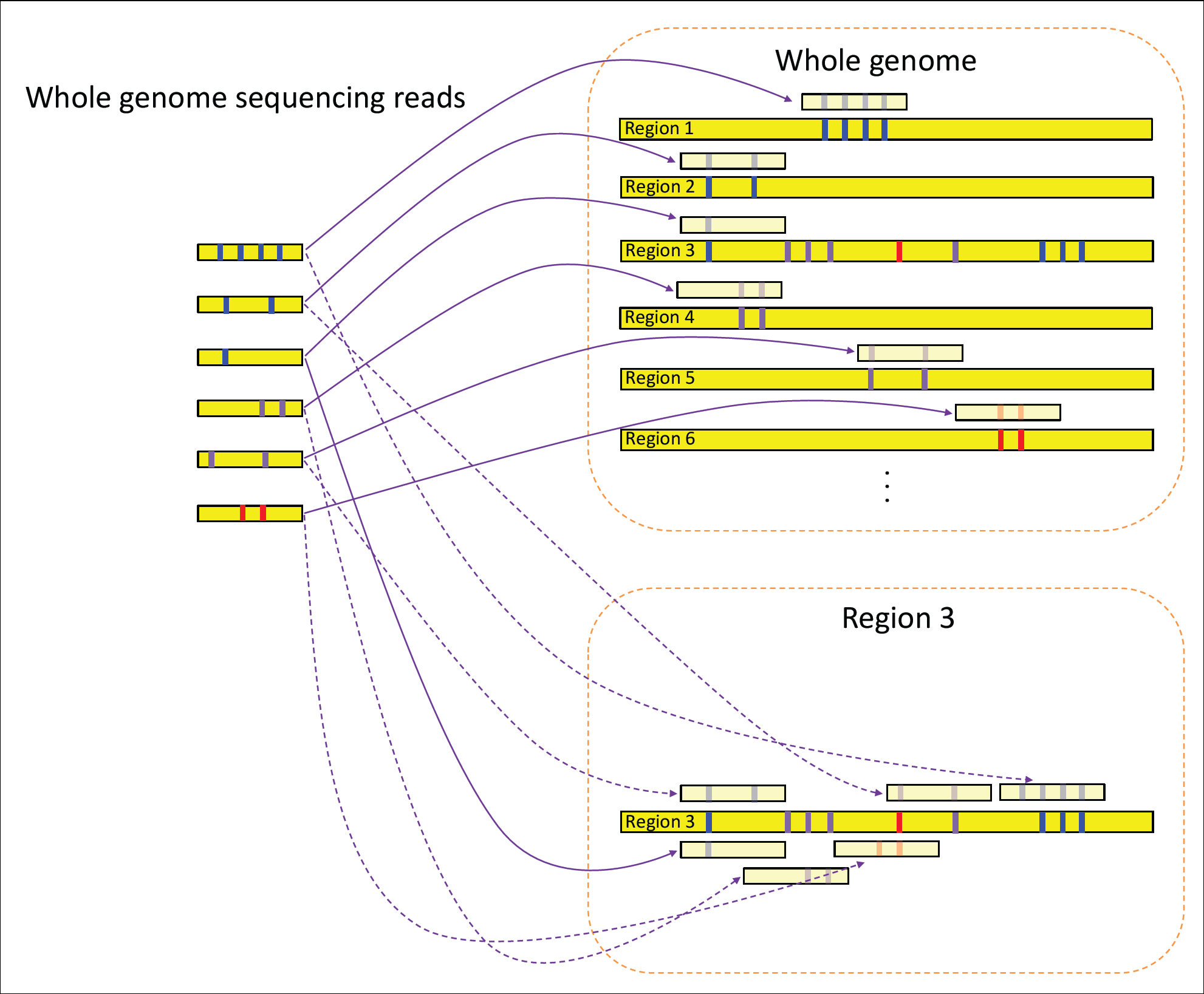
Incorrect reference leading to mis-alignment and bias. An illustration of the benefits of using the right reference/index when working with sequencing reads. The figure shows the alignment of reads to the whole genome (upper right) and to one particular genomic region denoted as Region 3 (lower right). When using the whole genome for aligning the six example reads, reads are perfectly aligned to the correct regions (regions 1, 2, 3, 4, 5, and 6). However, if the example reads are aligned using only one particular region (e.g. Region 3), five out of the six reads are incorrectly aligned to that region because alignment programs allow for a few mismatches. This excessive and loose alignment is a significant source of bias in analyzing genes or genomic regions. For example, in order to identify and extract reads that belong to the HLA-DRB1 gene from whole genome sequencing reads, one may attempt to align them only to the HLA-DRB1 gene region. In our experiment, we found that this strategy produced 1100 times more reads mapped to HLA-DRB1, compared to the alignments we obtained using the whole genome for alignment and extraction (HLA-DRB1 only vs. whole genome: 51844x vs. 46.1x coverage on NA12878, the latter being close to the expected 53.4x coverage). Thus, alignment accuracy can be greatly enhanced by using the right reference.

Once reads are extracted that belong to a particular gene or genomic region using a *Genotype* genome, HISAT-genotype performs further downstream analyses based on the read alignments: (1) typing and (2) gene assembly.

Typing is the process of identifying the two alleles (or the one allele if homozygous) for a particular gene that best match a given sequencing data set as shown in Fig 13.

**Fig 13.**
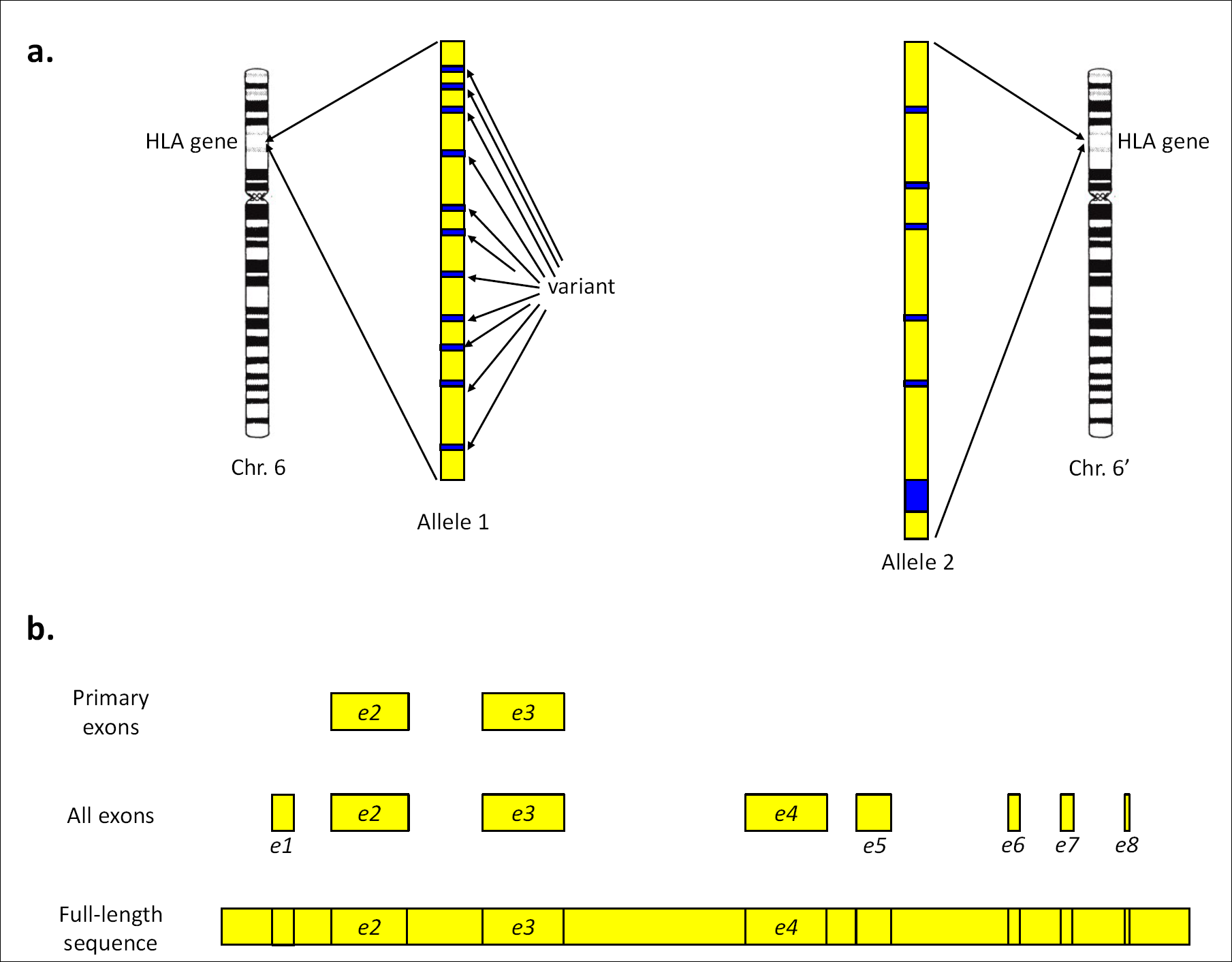
Two alleles of an HLA gene. *a*. Each person has two versions of a gene or a genomic region (e.g. HLA-A gene), shown in the figure in yellow with variants in blue, one version from the mother and the other from the father. *b*. The IMGT/HLA database includes many sequences for some key exons involved in core functions of proteins of HLA genes, but it contains far fewer complete sequences comprising all exons, introns, and UTRs of the genes. For example, 3,644 alleles have been classified so far for HLA-A. Although all alleles of HLA-A have known sequences for exon 2 and exon 3 (*e2* and *e3* in the figure), only 383 alleles have full-length sequences available. The sequences for the rest of the alleles (3,261 alleles) include either only all the exons (*e1* to *e8*) or a subset of them. For other HLA genes, HLA-B has 4,454 alleles, for 416 of which full sequences are available. HLA-C has 3,290 alleles, with only 590 fully sequenced, HLA-DQA1 has 76 alleles with 53 fully sequenced, HLA-DQB1 has 978 alleles with 69 fully sequenced, and HLA-DRB1 has 1,972 alleles, with only 43 fully sequenced. The chromosome 6 image is extracted from Fig 4-11 in *Molecular Biology of Cell* 5^th^ ed.

Because allele sequences are either only partially (just including exons) or fully (exons and introns) available, HISAT-genotype first identifies two alleles based on the sequences commonly available for all alleles, e.g. exons. During this step, HISAT-genotype first chooses representative alleles from groups of alleles that have the same exon sequences. Next it identifies alleles in the representative alleles that are highly likely present in a sequenced sample. Then the other alleles from the groups with the same exons as the representatives that remain candidates are included again for assessment during the next step. Second, HISAT-genotype further identifies candidate alleles based on both exons and introns. HISAT-genotype applies the following statistical model in each of the two steps to find maximum likelihood estimates of abundance through an Expectation-Maximization (EM) algorithm [24]. We previously implemented an EM solution in our Centrifuge system [25], and we integrated that solution into HISAT-genotype with modifications to the variable definitions as follows.

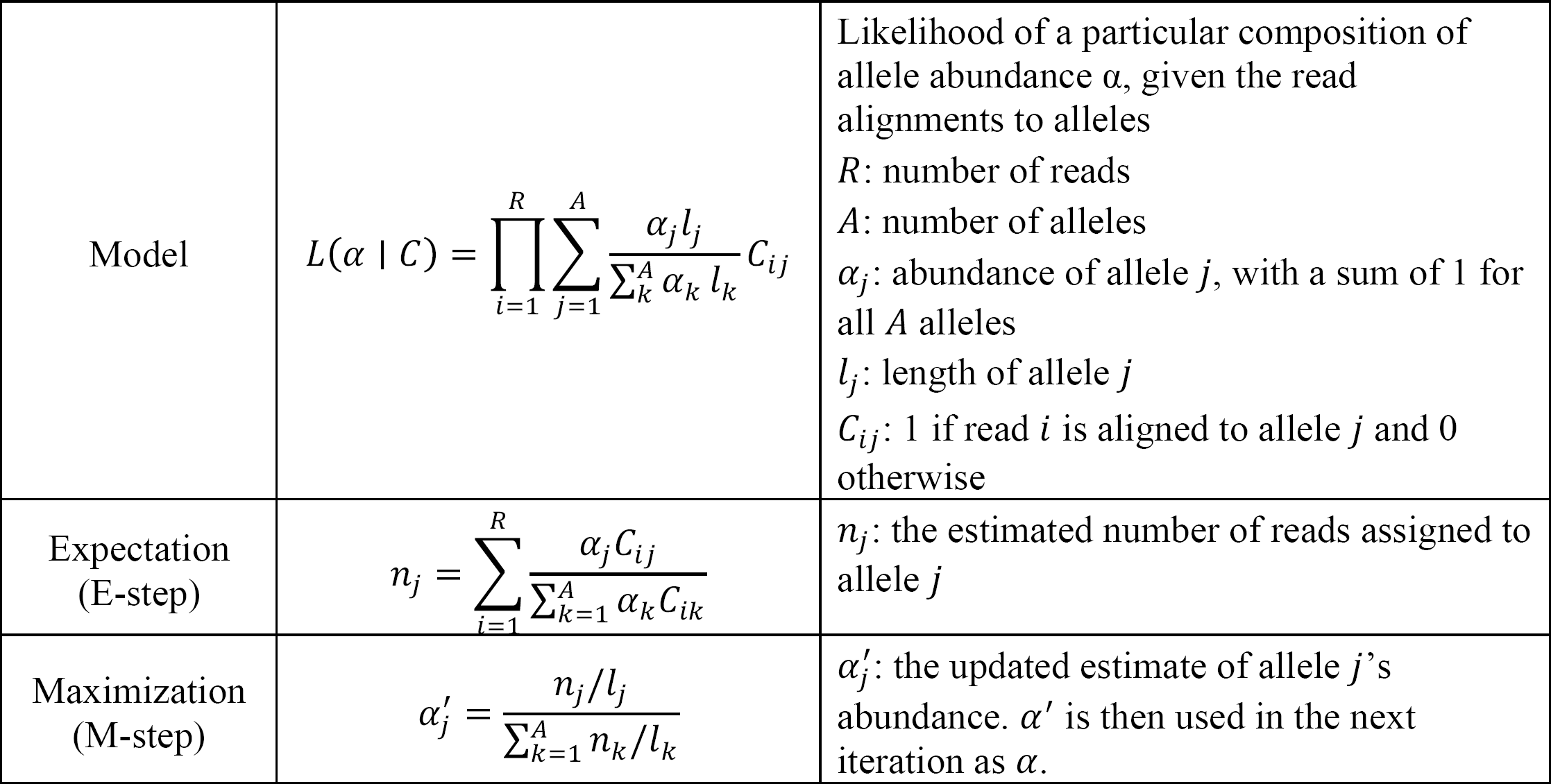

HISAT-genotype finds the abundances *α* that best reflect the given read alignments, that is, the abundances that maximize the likelihood function *L*(*α* | *C*) above by repeating the EM procedure no more than 1000 times or until the difference between the previous and current estimates of abundances, 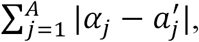 is less than 0.0001.

Here are some examples of typical abundances of alleles. If the sample has two alleles, a_1_ and a_2_, that exactly match the two alleles of the database, a_1_ and a_2_ are assigned abundances of approximately 0.5 each. If instead the sample is homozygous for that particular gene, the allele is assigned abundance of 1.0. If the sample has one allele (a_1_) exactly matching the database and the other (a_novel_) that does not perfectly match any allele but closely matches two alleles in the database (a_2_ and a_3_), we may ascertain an abundance of 0.5 for a_1_, 0.25 for a_2_, and 0.25 for a_3_. When paired-end reads of ≥100 bp with a sequencing depth of at least 30-50x coverage are used, HISAT-genotype is able to assemble full-length alleles and determine whether they are novel by comparing the assembled alleles with known alleles in the database, as described below.

Instead of directly assembling reads based on overlapping relations among reads (e.g. overlap-layout-consensus assembly approaches), HISAT-genotype splits aligned reads into fixed length segments called k-mers. These k-mers form an assembly graph (Fig 14) that enables the systematic assembly of alleles by handling noise and resolving assembly ambiguities.

**Fig 14.**
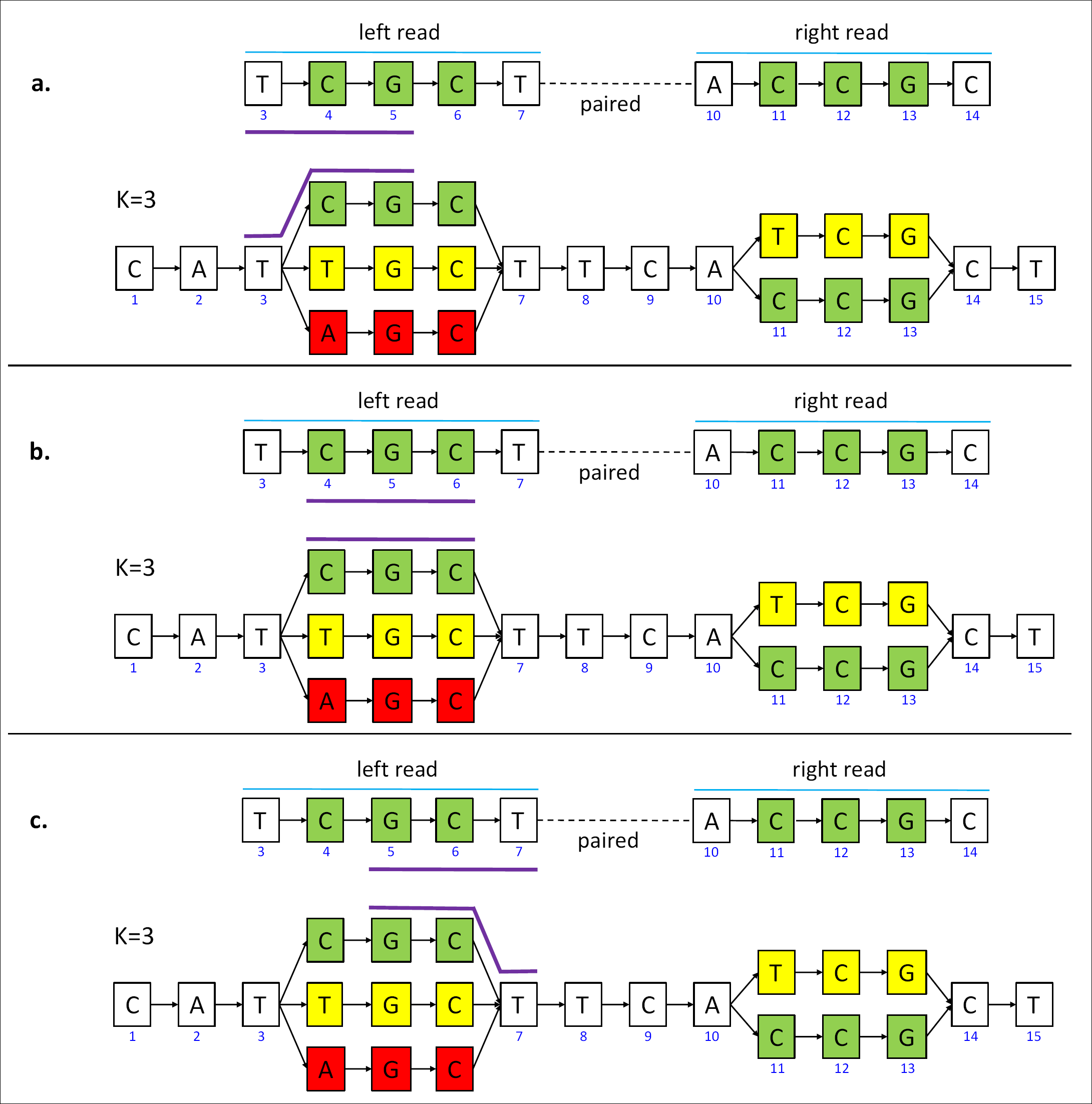
Guided k-mer assembly graph. HISAT-genotype splits aligned reads into fixed length segments called k-mers. For example, in the simplified case, reads are 5 nucleotides long and k is 3. A pair of reads are aligned at the 3rd location and the 10th location of the graph representation for the HLA gene, respectively. This example shows three k-mers from the *left* read alignments. This read consists of three k-mers, TCG, CGC, and GCT (the three k-mers from the right read would be similarly defined). Similarly, k-mers are extracted from other aligned reads (not shown here) and constitute the graph. When reads have the same k-mer, the graph has one path. When reads have different kmers, the graph has a corresponding number of branches. One path traversing the graph from left to right constitutes one potential allele sequence. This graph is here referred to as a guided k-mer assembly graph, with *guided* emphasizing that k-mers are placed according to their aligned locations.

As normal cells are expected to have two alleles if heterozygous or one allele if homozygous, one of the three k-mers in Fig 15a is likely to be from noise, meaning errors, generated during the sequencing or alignment stage. HISAT-genotype eliminates one of them by using the number of reads that support each k-mer as evidence. For example, if the k-mers shown in green and yellow are supported by 3 reads each, while the k-mer in red is only supported by one read, the program simply then removes the k-mer in red from the graph. After noise removal (Fig 15b), it is not yet clear which k-mers are linked to which k-mers from the same allele. For example, it remains undetermined whether CGC shown in green is connected to TCG in yellow or CCG in green. Pair information is then used to resolve this allele ambiguity. Suppose there are three pairs that support CGC and CCG in green, one of them like the example paired-end reads in the figure, and three pairs that support the k-mers in yellow. Drawing upon this pair-end read information, it can be determined that the k-mers shown in green come from the same allele, and similarly for the k-mers in yellow as illustrated in Fig 15c. In the case of two alleles having a large chunk of DNA sequence in common, fragments from which a pair of reads originate may not be long enough to resolve assembly ambiguity. For example, two known alleles A*01:01:01:01 and A*11:01:01:01 of NA12878 have the same ∼1,200 bp sequence in the middle while typical lengths of fragments range from 600 to 800 bps. In order to fully assembly alleles, HISAT-genotype makes use of alleles in the database to combine partial alleles into full-length alleles. This approach enables HISAT-genotype to assemble correctly all HLA-A alleles for the PG genomes, although this assembly can introduce bias toward alleles in the database. Due to many variants including insertions and deletions incorporated in the *Genotype* genome, it is often observed that a read can be locally aligned in multiple ways at approximately the same location as illustrated in Fig 16a in gray, where only one alignment is actually correct. If a program selects an incorrect local alignment, then that may in turn lead to choosing the wrong allele. HISAT-genotype handles such cases by choosing the most likely alignment using the aforementioned statistical model and EM method.

**Fig 15.**
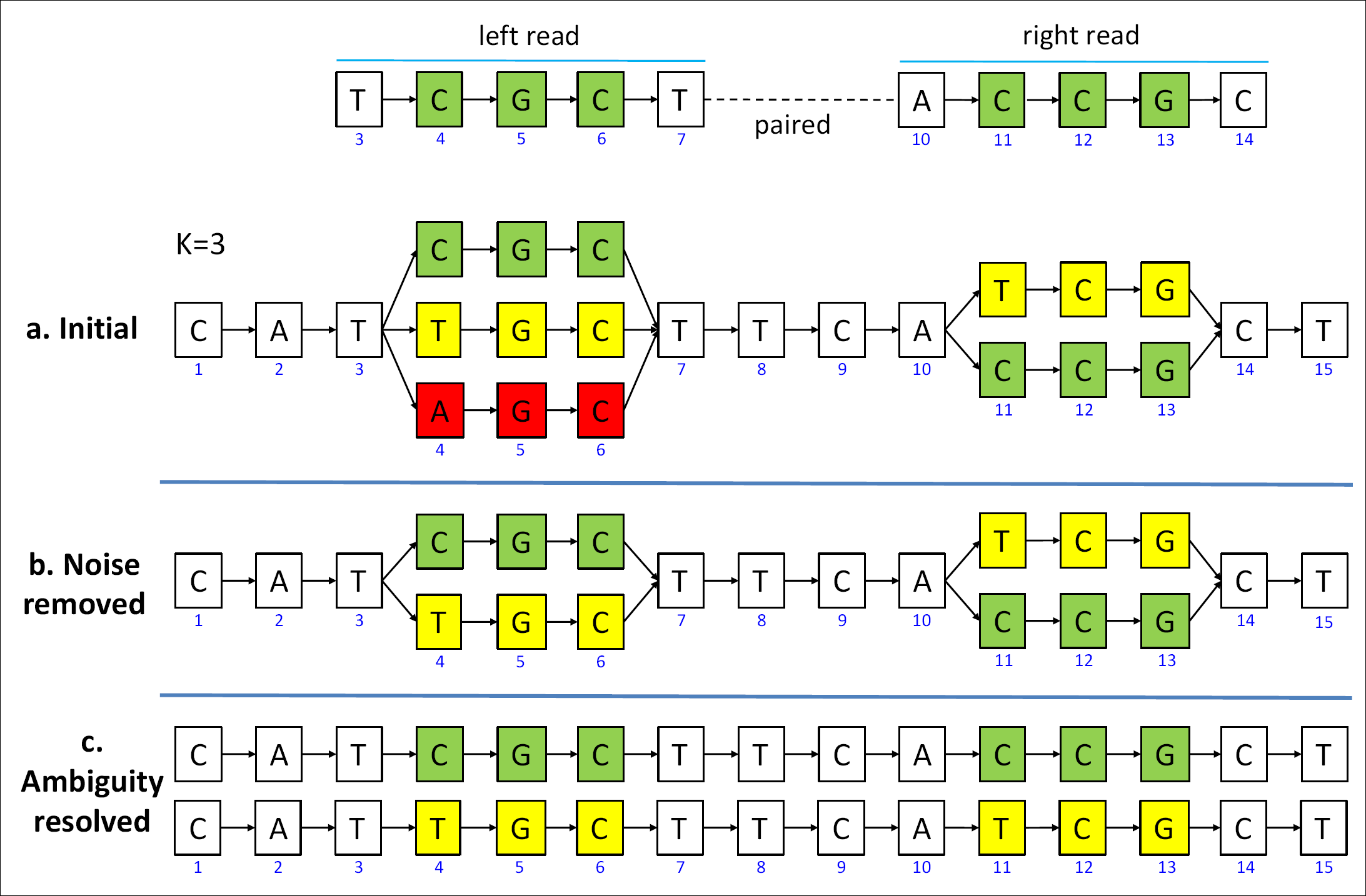
Assembly of two full-length alleles through guided k-mer assembly graph

**Fig 16.**
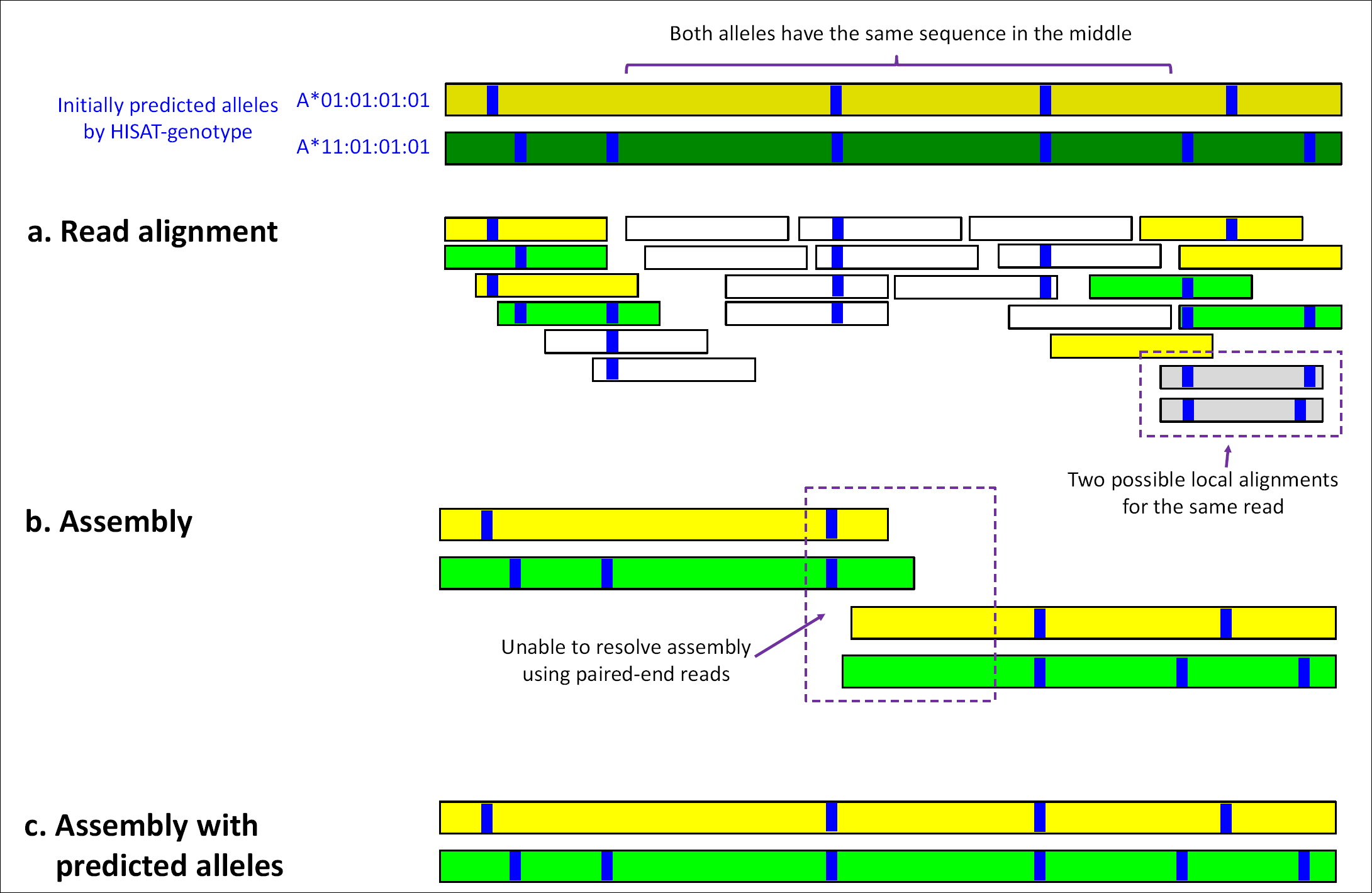
HISAT-genotype’s assembly output. The figure shows an abridged example of HISAT-genotype’s assembly output – see Supplementary Figure **4** for the full assembly output for NA12878 (one of the PG genomes) in PDF format. The first two bands are two alleles predicted by HISAT-genotype, in this case A*01:01:01:01 in dark green and A*11:01:01:01 in dark yellow. Each blue stripe indicates where there is a specific genomic variant with respect to the consensus sequence of the HLA-A gene. *a.* Depiction of shorter bands indicating read alignments whose color is determined according to their degree of compatibility with either the initially predicted alleles. If a read is equally compatible with both alleles, it is shown in white. Some reads can be locally aligned, i.e. aligned to virtually the same location with just different variants, such as when reads are aligned with or without deletions near their ends, in which case reads are displayed in gray. *b.* Since the two predicted (in fact true/known) alleles share a large common sequence, pair information is not enough to assemble full-length alleles, but only partial assemblies. *c*. So instead, making use of the predicted alleles enables the two full-length alleles to be assembled.

The algorithms described above are general enough to perform analysis of other regions of the human genome, as illustrated by HISAT-genotype’s accurate typing of 13 DNA fingerprinting loci plus the sex locus of the 17 platinum genomes.

## Acknowledgments

D.K. and S.L.S. performed the analysis and discussed the results of HISAT-genotype. D.K. implemented HISAT2 and HISAT-genotype. J.P. optimized the index building algorithm of HISAT2. D.K. performed wet-lab experiments. D.K. and S.L.S. wrote the manuscript. We would like to express our thanks to Kathleen Barnes and Michelle Daya for sharing Omixon’s HLA results with us. We would like to thank Ben Langmead and Jacob Pritt for their invaluable contributions to our discussions on HISAT2. We also greatly appreciate the generosity of Gaudenz Danuser and Dana Reed in providing wet-lab bench space and equipment for us. This work was supported in part by the National Human Genome Research Institute (NIH) under grants R01-HG006102 and R01-HG006677 to S.L.S and by the Cancer Prevention Research Institute of Texas (CPRIT) under grant RR170068 to D.K. All authors read and approved the final manuscript.

## Supplementary Figures

**Supplementary Figure 1.** CEPH pedigree #1463 consisting of 17 members across three generations

**Supplementary Figure 2.** HLA-A gene assembly of PG genome NA12892

**Supplementary Figure 3.** HLA-A gene assembly of CAAPA genome LP6005093-DNA_E03

**Supplementary Figure 4.** HLA-A gene assembly of PG genome NA12878

## Supplementary Files

**Supplementary File 1.** HISAT-genotype’s HLA typing results for 17 PG genomes on HLA-A, HLA-B, HLA-C, HLA-DQA1, HLA-DQB1, and HLA-DRB1

**Supplementary File 2.** HISAT-genotype’s HLA typing results for 917 CAAPA genomes on HLA-A, HLA-B, HLA-C, HLA-DQA1, HLA-DQB1, and HLA-DRB1. The CAAPA genome data is available from dbGaP as accession phs001123.v1.p1

**Supplementary File 3.** HISAT-genotype’s initial DNA fingerprinting results for 17 PG genomes

**Supplementary File 4.** PowerPlex^®^ Fusion results for 17 PG genomes (raw signal image data)

**Supplementary File 5.** List of alleles for 13 DNA fingerprinting loci and Amelogenin locus from NIST STR database

**Supplementary File 6.** List of 8 additionally incorporated alleles for 4 DNA fingerprinting loci D8S1179, D13S317, VWA, and D21S11

**Supplementary File 7.** HISAT-genotype’s DNA fingerprinting results for 17 PG genomes after incorporating 8 novel alleles from the 17 PG data

